# Single-cell resolution view of the transcriptional landscape of developing *Drosophila* eye

**DOI:** 10.1101/763243

**Authors:** Radoslaw Kamil Ejsmont, Grace Houser, Natalia Mora Garcia, Sara Fonseca Topp, Natalia Danda, Agnes Wong-Chung, Bassem A. Hassan

## Abstract

Faithful and reliable quantification of gene expression at a single-cell level is an outstanding challenge in developmental biology. Most existing approaches face a trade-off between the signal to noise ratio, resolution, and sensitivity. Here, we present a novel approach for in situ quantification of gene expression in a developing tissue. Our pipeline combines computational prediction of transcription factor targets, gene tagging, fluorescent reporter imaging, state-of-the-art image analysis, and automated cell-type identification. By applying this approach to identify the sequence of quantitative changes in gene expression which govern the development of the *Drosophila* neural retina, we demonstrate the feasibility of our method. We analyze the targets of Atonal (Ato), a transcription factor that controls the transition from eye disc progenitor cell to photoreceptor neurons. We utilized recombineering and genomic engineering to tag all predicted Ato targets with novel transcriptional reporters. These reporters enable following the expression of both regulator and regulated genes to accurately quantify their expression levels in individual cells. Our complete computational pipeline identifies nuclei in the eye discs and detects different states of cells as they progress through differentiation. Based on detailed gene expression analysis, our technique revealed genes likely to be direct Ato targets and provided insight into how gene expression changes drive the specification of photoreceptors.

## Introduction

Animal development is a complex process in which a single cell gives rise to a complex, multicellular organism. This process involves divisions, migration, differentiation, and death. Cellular differentiation is a multi-step process, where a cell faces consecutive decisions progressively defining its terminal fate. The defining transitions that a cell experiences, during development, are often regulated by key transcription factors. The complex genetic interactions, involved in cellular differentiation, require precise spatiotemporal control of gene expression. Gene expression can be directly quantified on both the protein and mRNA level. Protein-based methods include traditional western blotting and quantitative mass spectrometry that enables genome-wide analysis^1^. Measurements of gene activity are more commonly assayed on the messenger RNA level using quantitative PCR^2^, microarray analysis^3^, and next-generation sequencing^4^. Most direct methods of gene expression quantification require isolation of protein or mRNA from cells. This is laborious and results in loss of spatial and temporal resolution. The spatial resolution limitations of these methods can be overcome using the fluorescent in situ hybridization (FISH) technique. FISH can be applied to gather quantitative gene expression data at a single-cell resolution and below^5^. However, relative expression levels are difficult to compare between different genes due to varying affinities of in situ probes. Single-molecule fluorescent in situ hybridization (smFISH) addresses this issue by quantifying the number of objects (mRNA molecules), instead of the gross fluorescent signal per cell^6,7^. While smFISH provides absolute quantification of gene expression, it cannot be applied to living cells and requires very high-resolution imaging. Therefore this technique limits the number of cells in which mRNAs can be simultaneously quantified. The difficulties in the direct quantitative detection of mRNA in the developing cells led to the emergence of indirect methods that use various reporters as a proxy.

Indirect methods of gene expression quantification usually involve placing a reporter under the control of a gene’s promoter. This is followed by the detection of the reporter through biochemical assays, enzymatic reactions, or fluorescence. These methods mostly rely on tissue imaging and therefore provide very good spatial data. In genetic model organisms like *Drosophila* fruit flies, the worm *C. elegans* and zebrafish, the Gal4/UAS binary expression system was most commonly used to create enhancer traps^8^ and visualize gene expression. The expression of various reporter proteins, under the transcriptional control of the yeast upstream activating sequence recognized by Gal4, provided means for monitoring gene expression in a tissue-specific or temporarily triggered manner^9^. However, this method is not quantitative due to nonlinear signal amplification, caused by the Gal4 transcription factor. Another interesting method has been specifically developed to directly assay mRNA levels in living cells. A combination of the MS2 phage coat protein fused to a fluorescent protein and an mRNA carrying MS2 binding sites enables direct visualization of transcripts in cells and tissues^10,11^. While powerful, this technique is quantitative only in combination with single-molecule imaging, thus suffers from similar issues as smFISH. There is a clear window of opportunity for improvement of current collection and analysis of gene expression data by allowing researchers to collect larger, more meaningful datasets and to further enhance our understanding of animal development.

Work on the powerful model system *Drosophila melanogaster* has led to the discovery of many transcription factor families and their physiological functions. The formation of the crystalline neural retina in the fruit fly has long been used as a powerful model to study the genetic control of cellular differentiation. The fly neural retina consists of around 800 unit eyes or ommatidia, each containing 8 photoreceptor cells named R1-R8. The sequence of events in *Drosophila* retinal differentiation is controlled by three structurally and functionally conserved transcription factors called Eyeless/Pax6 (Ey), Atonal (Ato) and Senseless (Sens). The high functional conservation of these key transcription factors across animal species, suggests that observations obtained in *Drosophila* likely have relevance for the understanding of similar processes in mammalian systems.

Eyeless is the primary control switch for eye development. Ectopic expression of Ey in various epithelial primordia, the tissue in its earliest stage of development, leads to the formation of eye structures on *Drosophila* wings, legs, and antennae^12^. The functional role of *ey* in the control of eye development is conserved in the animal world^13^. While Eyeless triggers the development of the eye, Ato governs neuronal cell fate specification in multiple *Drosophila* sensory organs including olfactory^14^ and auditory^15^ organs, and the eye^16^. In the eye, Ato is required for the selection and specification of the first photoreceptor cell, called the R8, which later recruits the remaining seven photoreceptor neurons to form the ommatidium. Thus, in *ato* mutants, none of the photoreceptors differentiate and the neural retina fails to form. Ato belongs to the basic helix-loop-helix (bHLH) protein family. Together with the generally expressed bHLH cofactor Daughterless/E12 it forms a heterodimer and acts as a transcription factor. Target genes of Ato include *senseless* (*sens*), *fasciclin 2* (*Fas2*), *dacapo* (*dap*) and *Down syndrome cell adhesion molecule* (*Dscam*)^17^. *Senseless* is a target of Ato required for maintenance of the acquired neural cell fate. Sens together with another transcription factor called Rough (Ro) forms a bistable negative-feedback loop that allows the neural precursor cell to acquire and lock in its terminal R8 fate^18^. Therefore, these key transcription factors form the backbone of the eye development genetic program.

The genes described above, which together are key players of the photoreceptor specification network, have been identified either in genetic screens^19,20^, computationally^17^ or in chromatin immunoprecipitation (ChIP-chip / ChIP-seq) experiments^21^. While these methods enable the identification of players and their roles in this complex network, the data originating from them is largely limited. Genetic screens, powerful in that they provide *in vivo* data, are laborious and limited in the number of network members they reveal. Computational screens identify potential members of the network by searching for the enrichment of a motif, specific for a particular transcription factor in the genome. Target gene lists, produced computationally, contain both false-positives and false-negatives. The potential targets identified computationally, still need to be confirmed experimentally. The ChIP-chip and ChIP-seq experiments provide the most direct evidence of transcription factor binding, but they require a very large amount of material and high-quality antibodies for the analyzed transcription factors. Additionally, tissue-specific ChIP involves laborious and often disruptive sample preparation. Both ChIP-based and computational methods focus mainly on the identification of enhancer regions. However, it is difficult to extract if and how the overall expression of a potential target is influenced by the transcription factor. This is particularly important, as sometimes subtle quantitative spatio-temporal changes in gene expression can contribute significantly to cellular and tissue phenotypes.

Despite decades of research, only a basic and non-quantitative outline of the gene regulatory network promoting neural cell specification in the *Drosophila* retina is available. How this network is initiated and modified as cells transition through successive states of differentiation is essentially unknown. Equally unknown is how the activity of transcription factors acting as binary switches of cell fate, such as Ato, is translated into a spatiotemporally patterned expression of target genes. Here, we present a novel approach to quantify the expression of transcription factors and their targets. We combine computational target gene predictions with high-resolution imaging of genetically encoded reporters to collect single-cell resolution data in the developing retina. In this manuscript, we present the complete solution for single-cell gene expression quantification in whole tissue, the eye disc datasets we acquired, and insights into the Ato targetome.

## Results

### Selection of Ato target genes for imaging

We developed a novel pipeline (Fig. 1) to address issues with the insufficient resolution of classical gene expression data. The expression of Ato and its targets is captured at high dynamic range and resolution by quantitative imaging of the tagged alleles. Focusing on Ato, we computationally predicted 92 putative targets (including *ato* itself) in the eye disc using i-cisTarget^22^. For each predicted target, we identified available genomic clones from FlyFos (36) and p[ACMAN] (49) genomic libraries^23,24^ and tagged each with a T2A-Venus transcriptional reporter (Supplementary Fig. 1). All tagged fosmids and 10 BACs were selected for transgenesis (Supplementary Table 1). We prioritized transgenesis of target genes that are co-regulated by other members of eye and retina specification gene regulatory networks (6 carried in fosmid clones and 10 in BAC clones), namely by Ey and Sens. We successfully obtained transgenic lines for 38 Ato targets. The transgenesis for 8 tagged targets (7 FlyFos-based and 1 p[ACMAN]-based) failed to produce stable lines. In addition, we found that out of 38 lines, the integrity of 6 p[ACMAN]-based transgenes had been compromised during transgenesis. In total, clones for 32 target genes, including 7 genes co-regulated by *ey* or *sens*, were suitable for further analysis. For the visualization of Ato protein, we created a mCherry-tagged allele *in situ*, using the IMAGO technique^25^. The successfully obtained reporter transgenic lines were combined with the Ato[mCherry] line for imaging.

**Figure 1.**
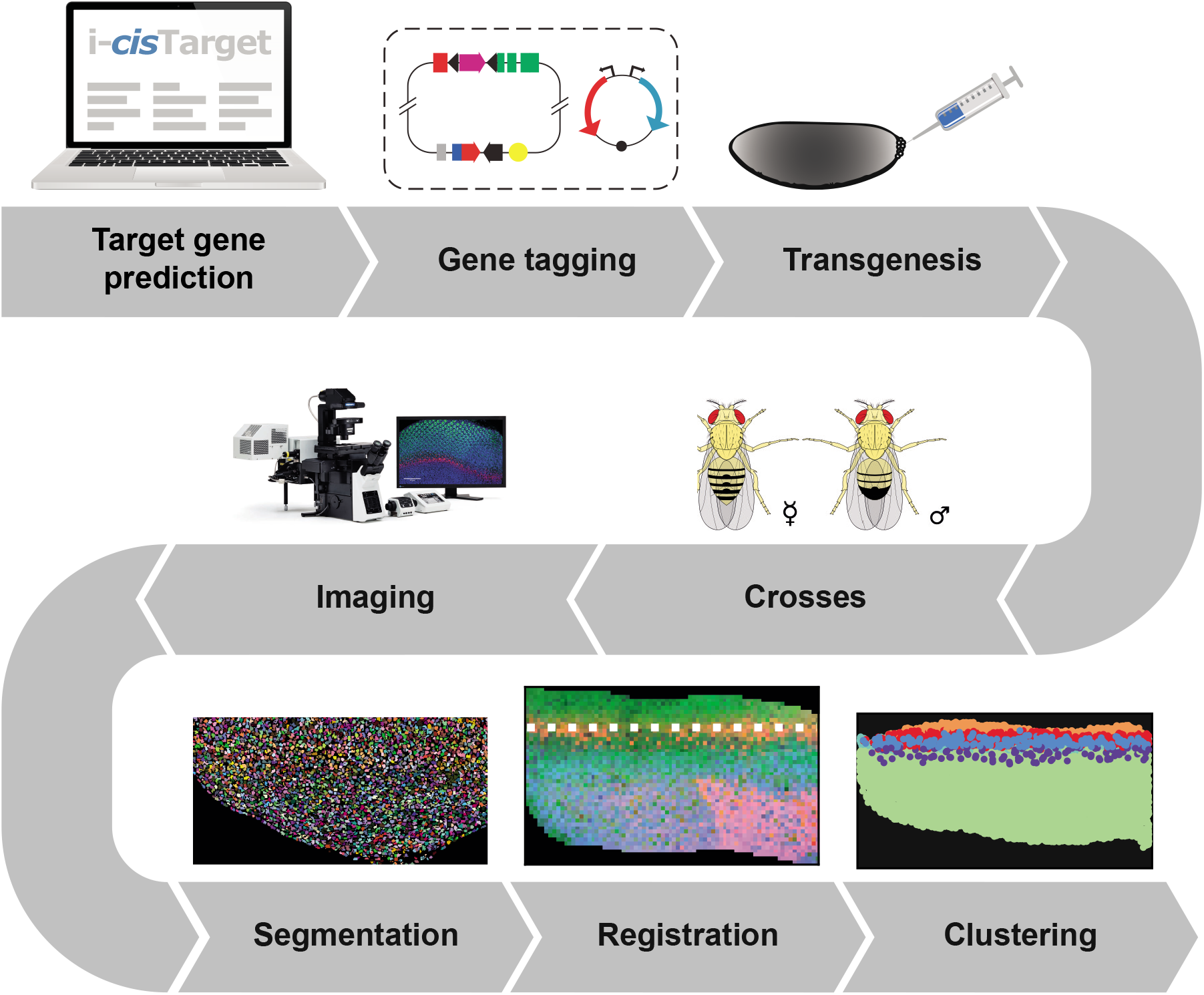
Experimental pipeline. Our experimental pipeline starts with computational prediction of putative Ato targets. Each target is tagged with a nuclear fluorescent reporter and the resulting tagged genomic constructs are injected into fly embryos. Tagged target genes are then combined with the fluorescently labeled Ato allele in a series of genetic crosses. The eye discs from third instar larvae are dissected and imaged. Nuclei from each image are segmented. Discs are aligned along the morphogenetic furrow, nuclear coordinates and signal intensities are normalized to the mean Ato intesity along the morphogenetic furrow. Nuclei are clustered based on their A-P position and the expression of Ato revealing different stages of R8 photoreceptor differentiation.

### Nuclei segmentation and registration

The first hurdle to single-cell quantification of gene expression in situ is the effective segmentation of individual cells in dense tissues. To overcome this hurdle, we developed an efficient nuclear segmentation algorithm (Supplementary Fig. 2). We imaged expression of Ato[mCherry] and 38 target gene transcriptional reporters in the eye discs dissected from wandering third instar larvae. We segmented individual nuclei from each confocal stack using DAPI staining as a nuclear marker. In further analysis, we represented each nucleus as a sphere with a volume equal to that measured from the original image. The signal intensity for each nucleus was measured as the mean photon count from the whole nuclear volume. Our segmentation algorithm sufficiently identified individual Ato-expressing R8 nuclei, despite being based solely on DAPI staining (Fig. 2ab). To facilitate image registration, we created a disc coordinate system. This system is based on the mean diameter of nuclei in each sample, their position relative to the disc edge, and the line of maximum Ato expression in the morphogenetic furrow (MF) (Fig. 2cd and Supplementary Fig. 3). From 389 imaged discs we obtained over 3.4 million nuclei. However, approximately 21% of nuclei were excluded from analysis because they had a volume deviating from the sample mean by more than 50%. This deviation was likely a result of over- or under-segmentation. The mean nuclear volume of ~30μm^3^, calculated from our data (Fig. 2e), is on the same order of magnitude as those measured by others^26,27^. The nuclear volume was relatively consistent across imaged discs, with the sample mean ranging from 22-38μm^3^ (Fig. 2f). The number of nuclei in the discs (Fig. 2g) varied between 5 and 17 thousand, depending on the disc age, and was consistent with the number of nuclei in the developing fly retina^28^. Finally, we measured the mean Ato[mCherry] signal intensity in the nuclei along the MF for each sample (Fig. 2h). The large variability (~90% of samples fall into values between 10 and 50) was expected. The observed variability is likely due to differences in fluorophore degradation during sample processing (dissection, mounting), as well as the time between sample mounting and imaging (ranging from 1 to 15 hours). To account for differences in fluorescence intensity between samples, we normalized measured intensities of Ato[mCherry] and the T2A-Venus transcriptional reporters to the mean intensity of Ato[mCherry] along the MF. For each nucleus, we computed the following parameters: xyz position in disc coordinate system; normalized intensities of Ato[mCherry], T2A-Venus and DAPI; and the prominence of Ato[mCherry] and T2A-Venus intensities. We define the expression prominence as a ratio between signal intensity for a particular nucleus, and that of its 26 nearest neighbors. In summary, we obtained putative Ato target gene expression data from almost 2.7 million segmented eye disc nuclei.

**Figure 2.**
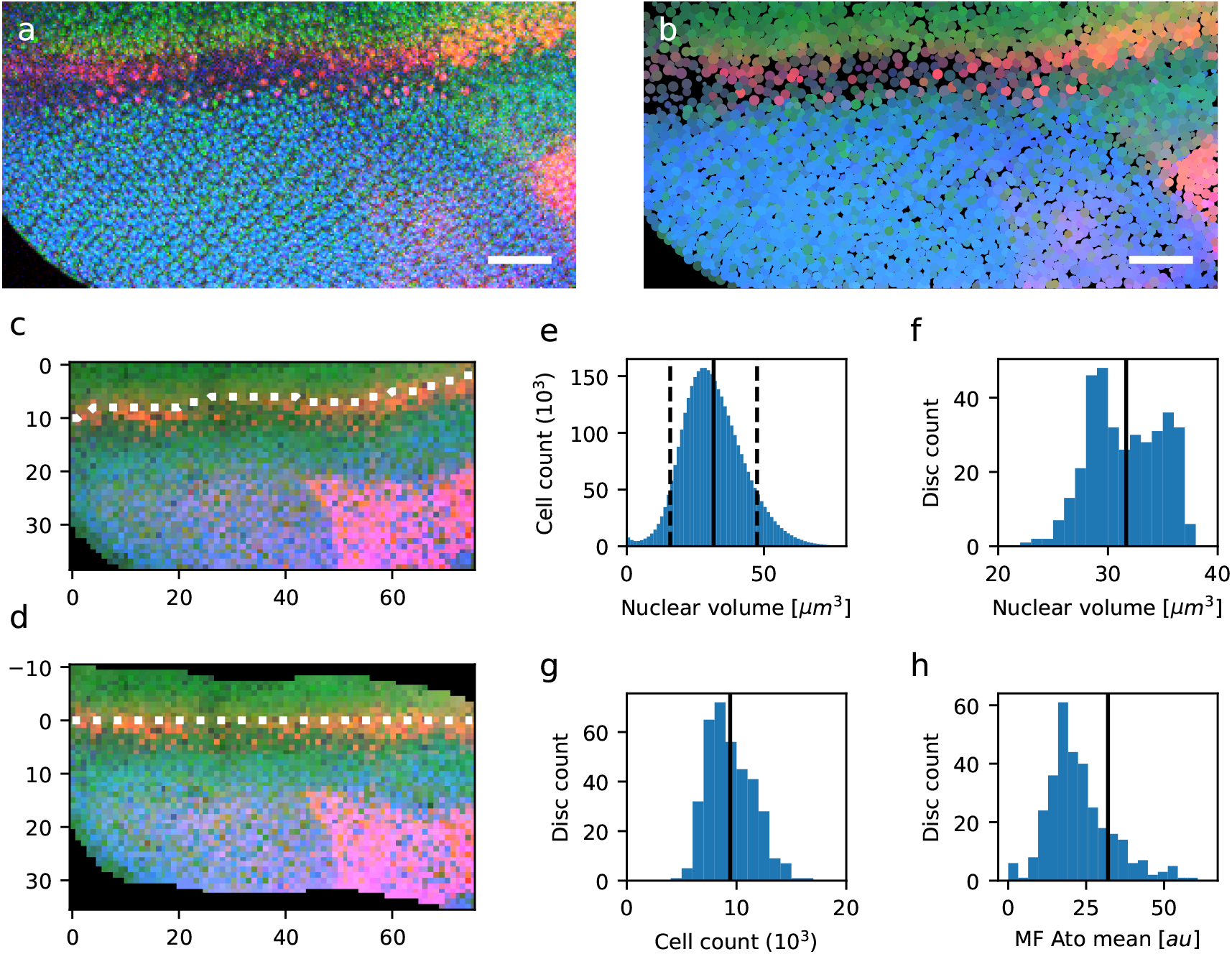
The quality of nuclear segmentation in the eye disc images. (**a**) Raw microscopic images were segmented into (**b**) nuclear point clouds. Both images show a single section (slice 40) of sample K21OU5. Nuclear DAPI staining is green, Ato[mCherry] fusion protein is red, *Lrch* (CG6860) transcriptional reporter is blue. The confocal image was acquired on the Olympus FV1200 point scanning confocal with a 40x/1.3 oil immersion objective. Scale bars are 30 μm. Anterior of the disc is at the top. (**c**) Segmented nuclei were projected (xyz mean-intensity projection) onto a 2-dimensional grid with normalized nucleus diameter (to the mean diameter, per sample) as a unit. Morphogenetic furrow (MF, dotted line) was detected as a line of maximum Ato expression in the anterior part of the disc. The vertical axis is the A-P distance from the anterior edge of the image. The horizontal axis is the D-V distance from the disc edge. (**d**) The vertical (y) coordinates of nuclei were transformed so that the MF forms a straight line at y=0. The y-axis is the A-P distance from the MF. The horizontal axis is the D-V distance from the disc edge. (**e**) Distribution of nuclear volumes across all imaged samples. The solid line indicates the mean value. Dashed lines indicate 50% difference from the mean. (**f**) Distribution of mean nuclear volumes per sample. The solid line indicates the mean value. (**g**) Distribution of total cell count per sample. The solid line indicates the mean value. (**h**) Distribution of the mean Ato[mCherry] signal intensities along the MF per sample. The solid line indicates the mean value.

### Expression of Ato and its targets in the eye disc

With single nuclei data in hand, we sought to understand the relationships between Ato expression and that of its potential targets. To this end, we combined Ato[mCherry] expression data from all samples and found that our tagged protein reporter recapitulates Ato expression pattern very well. Ato is first expressed by most cells (except the peripodial membrane) in a narrow band along the MF (Fig. 3a), later however its expression gets resolved only to R8 photoreceptors posterior to the MF (Fig. 3b). The expression of the *ato* transcriptional reporter follows a similar pattern and is higher in the R8s. However, it’s detectable much further posterior to the furrow (Fig. 3ce), especially in the R8 photoreceptors (Supplementary Fig. 4), and the prominence of expression in the R8s is lower for mRNA than protein (Fig. 3bd). The observed persistence of ato mRNA far beyond the MF supports the previously reported presence of a phosphorylated form of Ato in late R8s^29^. Along the D-V axis, we observed no significant variation in Ato levels, except for ~25% (protein) and ~33% (reporter) lower expression on the disc edges (Fig. 3f). In samples based on the FlyFos genomic clones we found, especially in the discs with high MF progression, strong expression of 3xP3-dsRed FlyFos selectable marker in the posterior of the disc, the optic nerve, and on the disc surface. This signal did not overlap with the expression domain of Ato and was easily distinguishable from the Ato[mCherry] signal (Supplementary Fig. 4). In accordance with these observations, both the Ato[mCherry] allele and the *ato* transcriptional reporter recapitulated the known features of Ato expression.

**Figure 3.**
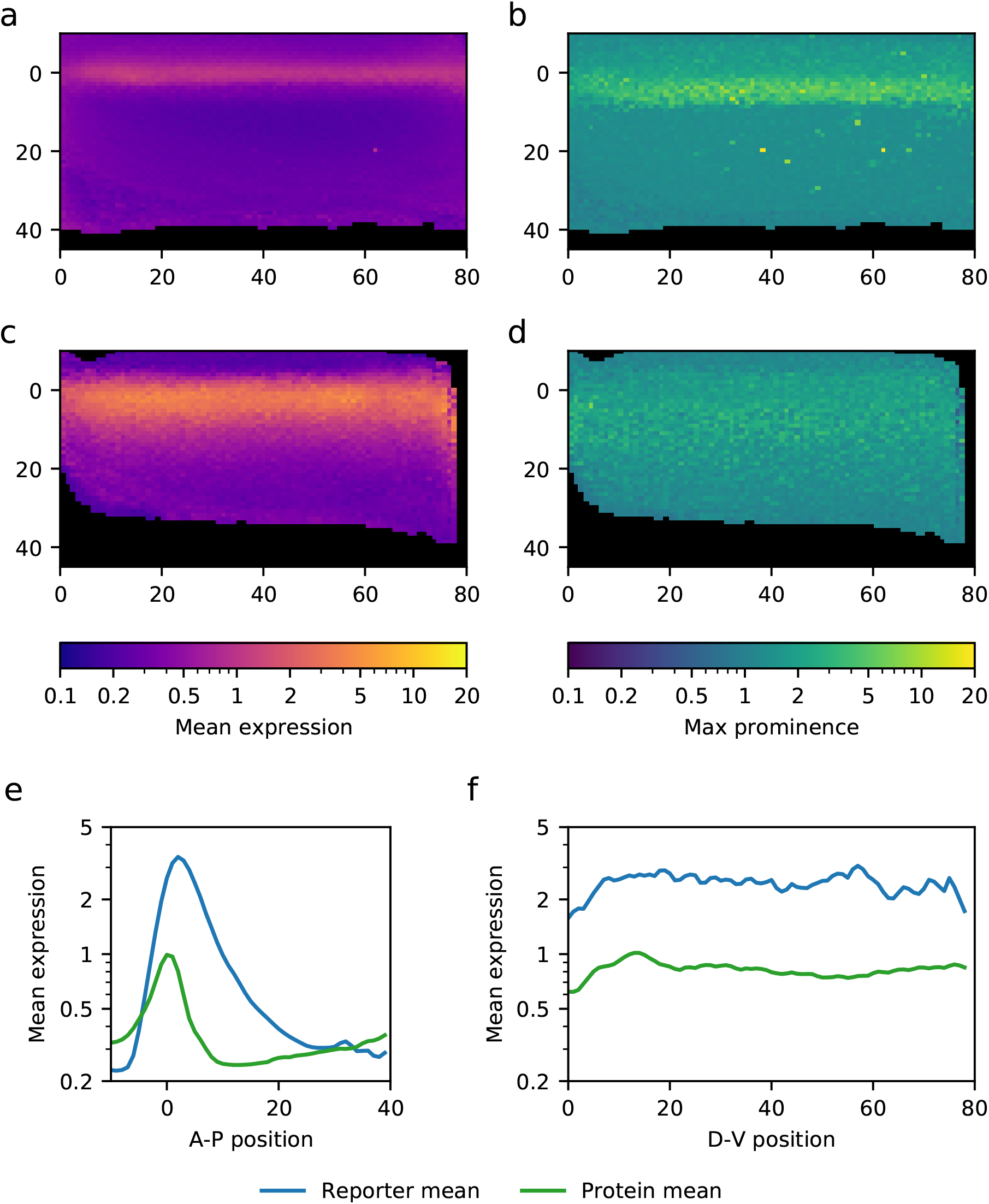
Expression of Atonal protein and mRNA in the eye disc. (**a**) xyz mean intensity projection of normalized Ato[mCherry] protein fusion expression from 55 p[ACMAN] BAC samples. The vertical axis is the A-P distance from the MF. The horizontal axis is the D-V distance from the disc edge. Axis unit is the normalized nuclear diameter. (**b**) xyz maximum projection of Ato[mCherry] prominence in the same samples. (**c**) xyz mean intensity projection of normalized ato-T2A-Venus-NLS transcriptional reporter expression. (**d**) xyz maximum projection of the transcriptional reporter prominence in the same samples. (**e**) Anterior-posterior (A-P) gene expression profile of Ato protein (green) and the *ato* transcriptional reporter (blue). Protein samples the same as in (a, b), transcriptional reporter samples the same as in (c, d). (**f**) Dorsal-ventral (D-V) gene expression profile of Ato protein (green) and the *ato* transcriptional reporter (blue).

We captured expression data of comparable detail for the predicted Ato target genes (Supplementary Fig. 5). Most (27) genes were expressed in the eye disc. The expressed genes fall into four spatially defined categories (Supplementary Table 2). The first category contains 12 genes whose expression starts within the MF. This group is highly enriched (6/12) for genes involved in the Notch signaling pathway. The nine genes in the second category are expressed immediately posterior to the MF. The four genes that belong to the third category are expressed posterior to the MF. The last category comprises two genes expressed both anterior and posterior to the MF (*dacapo, SRPK*). We saw almost ubiquitous expression of *dacapo* (*dap*) reporter, which is surprising as its expression was previously well described^30^ to be specific to R2/R5 precursors. We suspect that the upstream sequence of the fosmid carrying the *dap* reporter (~3kb) is insufficient to recapitulate the native expression pattern, and therefore we excluded *dap* from the further analysis. The expression of five target genes was not detected, including *sanpodo* (*spdo*), *phyllopod* (*phyl*), *CG31176, CG17378*, and *CG30343*. Lack of the expression of *phyl* is clearly contradicted by others^31,32^, indicating either damage to the fosmid carrying the reporter or the lack of regulatory elements necessary for eye disc expression of *phyl* in that fosmid. The gross analysis of putative Ato targets allowed us to identify which genes are more likely to be regulated by Ato. This includes genes that are expressed within, or in the proximity of the MF. However, we found that a closer look at how the expression varies across different cell types is essential to attribute weather these genes are direct Ato targets.

### Classification of cell types during R8 specification

We asked whether the quantity and quality of our dataset would allow the classification of cell state transitions during differentiation. Having detailed single-cell resolution Ato expression data from hundreds of samples, we classified cells based on their position along the anterior-posterior (A-P) axis, Ato expression level, and its prominence. We expected to primarily identify at least four classes of cells: cells anterior to the furrow (Pre-MF), Ato-expressing cells in the furrow (MF-medium), Ato-expressing R8 photoreceptors posterior to the MF (R8) and the remaining cells posterior to the furrow (post-MF). We hypothesized that the differences in Ato expression and prominence in these classes would be sufficient for clustering. Surprisingly our clustering algorithm failed to identify these classes (classifying some cells in the MF as cells anterior to the MF or as R8s) unless the number of clusters was increased to six (Fig. 4ab). The two additional clusters contained MF nuclei that do not express Ato (MF-low) and MF cells with high Ato expression and prominence (MF-high). The former class contains the peripodial membrane cells, while the latter we have attributed to the Ato-positive cells forming intermediate and equivalence groups. The A-P extent of each cluster is summarized in Supplementary Table 3. The expression level of Ato is the highest in the MF-high class, followed by the R8 and the MF-medium classes. The prominence of Ato expression is the highest in the R8 class, followed by the MF-high class (Fig. 4c). Mean position, Ato expression, and prominence in each class are consistent across all analyzed samples, with the largest variability in the pre-MF and post-MF classes (Fig. 4d). The mean number of segmented nuclei in each class varies between 5904±1517 (69%) in the post-MF class, 1322±831 (16%) in the pre-MF, 623±209 (7%) in the MF-medium, 378±172 (5%) in the MF-Low, 189±61 (2%) in MF-High and 92±23 (1%) in the R8 classes (Fig. 4e). The expression of Ato in the R8 remains relatively high up to six rows posterior to the MF, therefore yielding ~15 Ato-positive R8 photoreceptors per row of cells. Given that R8 cells appear every second row in our coordinate system, the number of R8s selected during one cycle is approximately 30, which is consistent with the literature^28^ and manual examination of our images. Consequently, the number of cells in the intermediate and equivalence groups should equal ~450 cells versus an average of 189 cells that we attributed to the MF-high class. Some cells from intermediate and equivalence groups were therefore assigned to the MF-medium class due to lower Ato levels or lower Ato prominence, suggesting high variability of the Ato protein levels during the initial steps of R8 specification. With an automated cell type classification system, we could proceed to profile how the expression of putative Ato targets changes during the R8 specification.

**Figure 4.**
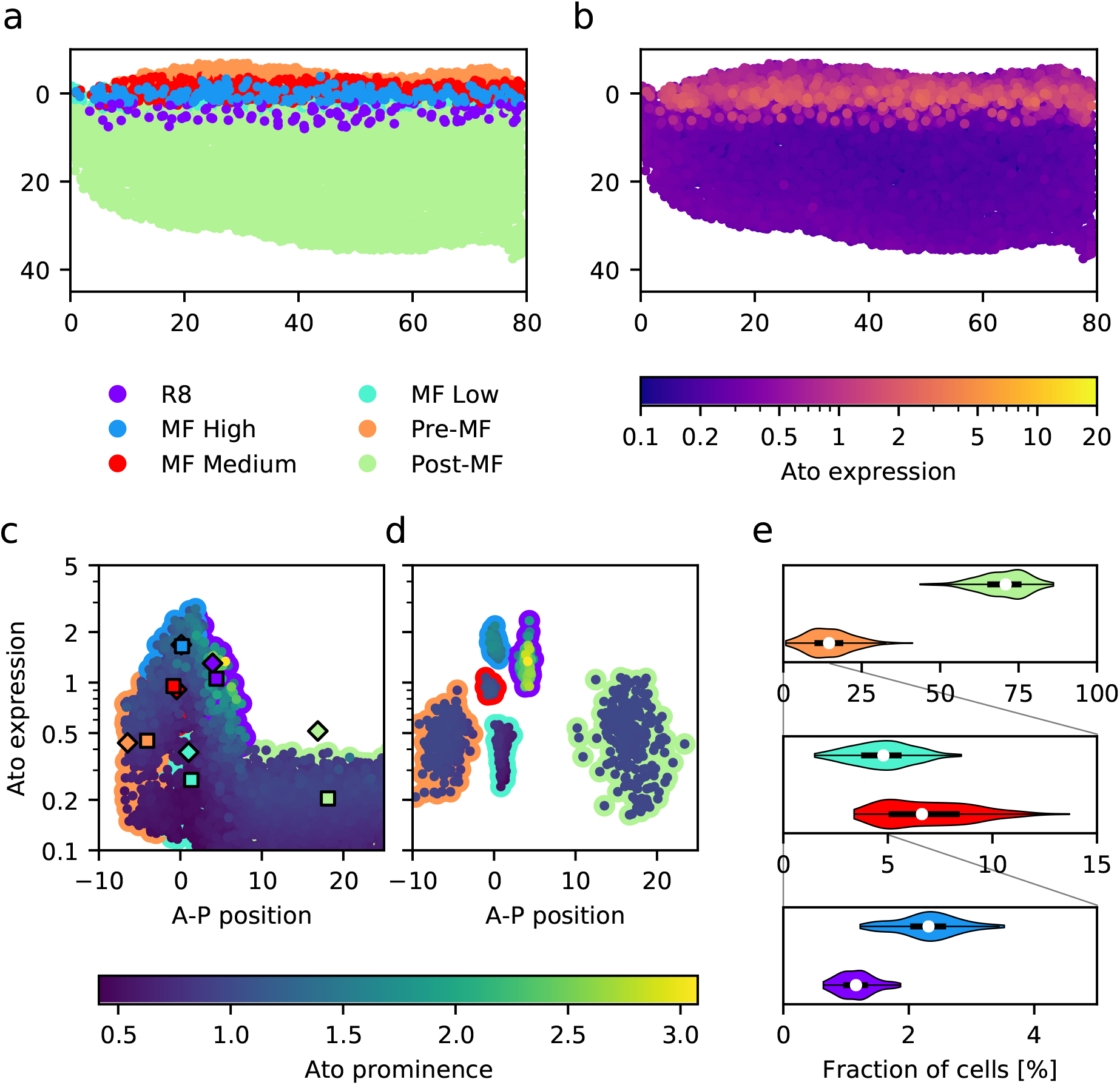
Cell type classification in the eye disc. (**a**) color-coded result of cell-type classification based on Ato[mCherry] expression in the eye disc (violet - R8, blue - MF-High, red - MF-Medium, cyan - MF-Low, orange - Pre-MF, green - Post-MF). Single disc, sample O4UW6B. The vertical axis is the A-P distance from the MF. The horizontal axis is the D-V distance from the disc edge. Axis unit is the normalized nuclear diameter. (**b**) Ato[mCherry] expression in the same sample. (**c**) Ato[mCherry] expression (vertical axis) and prominence (inner color) in different cell classes (outer color) along the A-P axis. These three variables were used for cell-type classification. Colored squares represent the cluster centroids in the sample O4UW6B. Colored diamonds represent global cluster centroids calculated from all samples. (**d**) Cluster centroids from every analyzed sample. (**e**) The fraction of cells that belong to each cluster, distribution across all analyzed samples.

### Expression of predicted Ato targets during R8 specification

To identify which of the predicted Ato targets have expression profiles related to Ato, we examined their expression levels in different cell classes and their expression profiles along the A-P axis. We found four main profiles that the predicted genes followed (Fig. 5a). Ten genes (Supplementary Fig. 6: *ato - seq*) follow the levels of Ato strongly, with the expression rising between MF-medium, MF-high and R8 classes. Expression of eight of these genes is the highest in the Ato-positive R8 cells, the remaining two genes (*Brd* and *E*(*spl*)*mδ-HLH*) are expressed more in the Ato-negative cells immediately posterior to the furrow. Two additional genes (Supplementary Fig. 6: *CG13928, Lrch*) also follow Ato levels, however to a much lesser extent, and with much shallower expression gradient within the MF. The *SR Protein Kinase* (Supplementary Fig. 6: *SRPK*) is expressed at high levels throughout the disc, though higher in the MF-high and the R8 cells. Among the genes that do not clearly respond to the varying Ato levels, the expression of three (Supplementary Fig. 6: *βTub60D, rau*, and *scrt*) onsets in MF proximity, with the highest levels in Ato-negative cells. Four genes (Supplementary Fig. 6: *Abl, CG17724, CG32150, DAAM*) do not exhibit differential expression in the Ato-positive cells. Five genes (Supplementary Fig. 6: *CG15097 - nSyb*) were not expressed in the proximity of the furrow.

**Figure 5.**
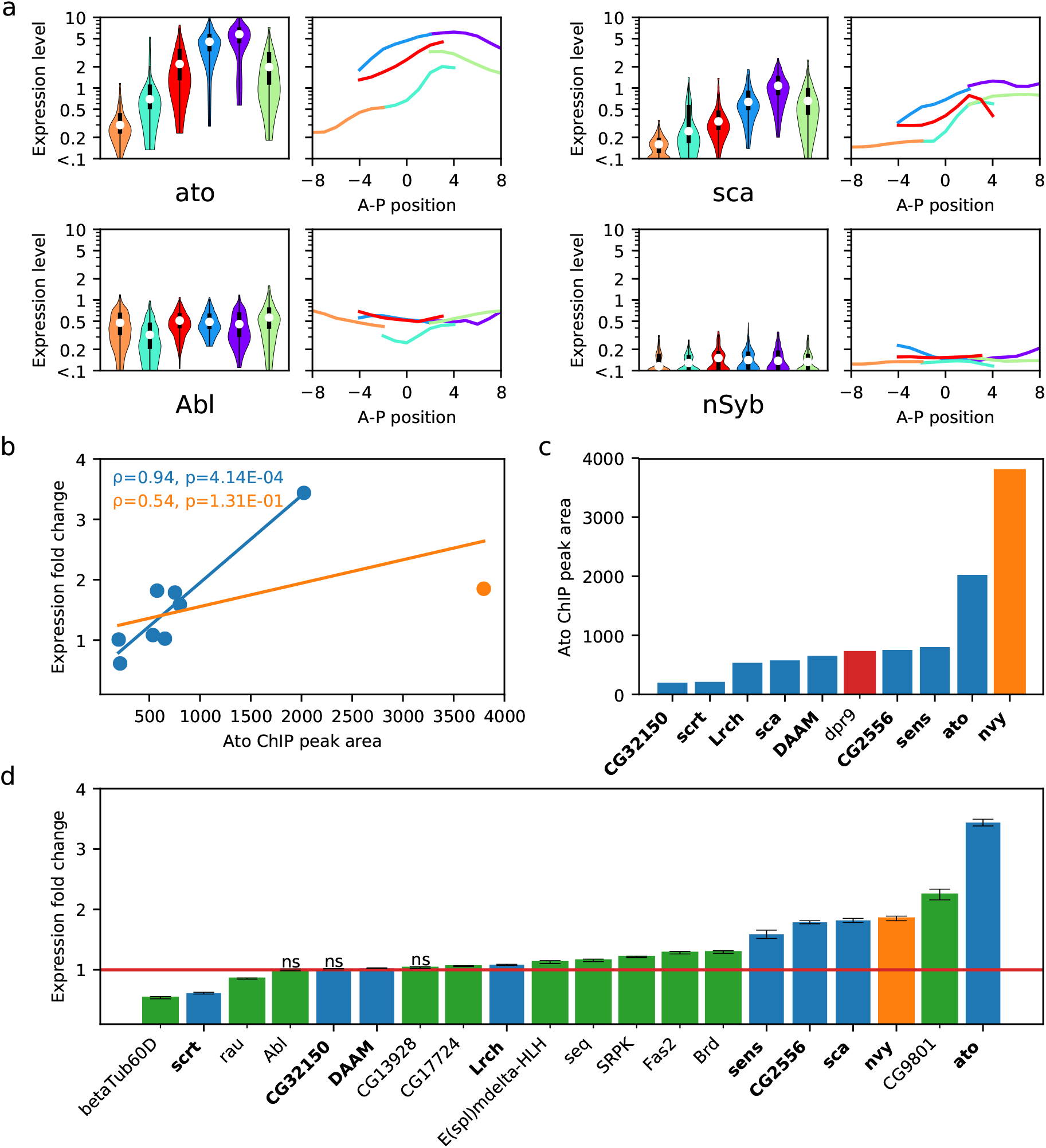
Expression of the putative Ato targets in different cell types. (**a**) Distribution of the putative target gene expression levels in different cell types in the vicinity of the morphogenetic furrow (violin plot) and the A-P expression profiles in each cell type (line plot). Color code is the same as in figure 4 (violet - R8, blue - MF-High, red - MF-Medium, cyan - MF-Low, orange - Pre-MF, green - Post-MF). (**b**) Expression fold change between Ato-high (MF-High and R8 classes) and Ato-low cells (MF-Low, Pre-MF, and Post-MF) plotted against Ato ChIP-seq peak area. The outlier (*nvy*) is plotted in orange, the remaining genes represented in both datasets are plotted in blue. The lines show the linear least-squares regression with the outlier included (orange) and excluded (blue). (**c**) ChIP peak area for genes included in this study. One that was not expressed in the MF vicinity in our imaging but has significant ChIP peak (*dpr9*) is plotted in red. The gene with disproportionately high ChIP peak (*nvy*) is plotted in orange. The remaining genes are plotted in blue. Genes represented in both ChIP and imaging datasets have their names printed in bold. (**d**) Expression fold change between Ato-high and Ato-low cells. Genes without ChIP peaks were plotted in green. The gene with disproportionately high ChIP peak (*nvy*) is plotted in orange. The remaining genes are plotted in blue. Genes represented in both ChIP and imaging datasets have their names printed in bold. Error bars represent propagated standard error of the mean (SEM).

With detailed high resolution expression data in hand, we asked how well expression profiles of Ato target genes correlate with Ato binding to target gene enhancers. To this end, we performed Ato ChIP-seq in the eye discs. We compared the upregulation of these genes in cells with high (MF-high, R8) and low (Pre-MF, MF-Low, Post-MF) Ato levels, to the Ato binding data from ChIP-seq (Fig. 5bcd). Ato binds strongly enhancers of two (*ato, nvy*) out of six most upregulated genes (*ato, CG9801, nvy, sca, CG2556, sens*), three are bound moderately (*sens, CG2556, sca*). Exceptionally strong binding of *nervy* (*nvy*) by Ato is not reflected in our expression data. Enhancers of *Lrch*, a weakly upregulated gene, were moderately bound by Ato. The binding of *CG9801* enhancers was not supported by ChIP-seq data. Interestingly, we found ChIP peaks for genes not expressed in the MF (*dpr9*), or those that appear to be invariant to the Ato levels (*CG32150, DAAM, scrt*). Surprisingly, we did not find ChIP-seq peaks for six genes upregulated in Ato-positive cells. Keeping in mind the lower cellular resolution of ChIP data, we find an overall strong correlation (ρ=0.94, p=4.14E-04, excluding nvy as an outlier) between the degree of binding in ChIP experiments and gene expression regulation in our imaging datasets. Based on the expression data in different classes and the enrichment of *Ato* binding sites in the gene vicinity (based on the i-cisTarget analysis), we propose that 13 out of 31 analyzed genes (*ato, Brd, CG2556, CG9801, E(spl)mδ-HLH, Fas2, nvy, sca, sens, seq, CG13928, Lrch*, and *SRPK*) are immediate and direct targets of *Ato* in the eye disc. Our data on five genes (*CG31176, DAAM, dila, rau*, and *spdo*) contradicts previous predictions^17^, as we did not find sufficient evidence supporting their direct regulation by Ato. However, as the expression levels of these genes in the MF area was close to our detection threshold (0.2 normalized units), we can neither confirm nor exclude these genes as Ato targets.

## Discussion

Methods to study developmental processes at a single cell level have long been a subject of intense technology development. Single-cell RNA sequencing in combination with clustering and data mining tools, such as SCENIC^33^ and SCope^34^ enable identification of cell types and states that cells transition through during differentiation. scRNA-seq, while providing a quantitative whole-transcriptome view at the level of individual cells, yields data of limited dynamic range and low signal to noise ratio^35^. Complex sample preparation and the desire to maximize sequencing depth limit the number of analyzed cells in most developmental samples from *Drosophila* to several thousand (rarely exceeding 20k). As a consequence, the resolution of cell-type identification using scRNAseq is low. Mapping scRNA-seq data onto FISH datasets^36,37^ introduces spatial dimensions to the single-cell transcriptional analysis. However, due to limitations of the scRNAseq, it enables only a rough estimation of the tissue regions that the individual cells originate from.

Our approach enables large scale quantitative gene expression analysis of a transcription factor targetome at single-cell resolution. This approach works on whole-tissue level, with very high sensitivity and dynamic range, while preserving spatial information. Thanks to whole tissue imaging, we are able to gather gene expression data from millions of cells. We have both spatial information as well as the precise measurements of expression levels for the transcription factor driving the studied developmental process. Thus, our approach enables us to identify transient states that cells progress through during differentiation. Unlike in approaches relying on vast amounts of noisy full transcriptome data, we find that precise analysis of the expression of a single key gene is sufficient to identify these states. Together with cell-type identification, our disc coordinate system (Supplementary Fig. 3) enabled identification of cells with the same properties in different samples, and thus to assert expression levels for all assayed genes during differentiation. As differentiation in the fly retina progresses as a wavefront along the A-P axis, the distance from the MF defines the developmental age of each cell. With expression data on both the transcription factor and its putative target genes, we are therefore able to find a spatial and temporal relationship between the levels of Ato and the expression of its target genes.

Based on this relationship, we were able to identify 13 genes as likely direct Ato targets. Five of these genes (*ato, E(Spl), Fas2, sca, sens*) were previously identified by others using both computational approaches and enhancer reporter assays^17^. We provided supporting evidence for four genes (*CG2556, CG9801, nvy, SRPK*) that were previously only predicted computationally^17^ and identified three new targets (*Brd, seq, CG13928*). Interestingly, none of the direct Ato targets we identified here encode for structural or neurofunctional proteins, but rather for members of the Notch or EGFR signaling pathways and other regulators of gene expression. This suggests that Ato regulates only the switch of cell fate, and the downstream differentiation program is executed through changes in the signaling state of the cell and fine-tuned through specific transcription factors such as *sens* and *seq*.

While the method presented here has been specifically tailored to the analysis of gene expression in the developing Drosophila retina, it is generalizable. The expression of any developmental transcription factor can be used as a landmark, in a similar fashion to how we used the expression of Ato. Depending on how differentiation progresses in a tissue of interest, different coordinate systems can be created. In the mammalian neocortex, like the fly retina, cells are arranged in a spatially layered order reflecting temporal specification. However, in other system, like the inner proliferation center (IPC) of developing Drosophila brain^38^ for example, a radial coordinate system could be suitable. With the advances in tissue culture and imaging techniques, such as lightsheet microscopy^39^ and clearing techniques our approach could be implemented in deep tissues, such as the mammalian brain. Deep learning-based analysis^40,41^ could further improve the image segmentation and cell-type classification, which could help to better assess genes expressed at very low levels.

## Acknowledgements

We are grateful to Pavel Tomancak, Hugo Bellen and BACPAC Resources Center for the genomic clones. We thank Mihail Sarov for the Red/ET recombineering plasmid. We are grateful to Stein Aerts for help with i-cisTarget analysis. We thank the Bloomington stock centre for fly stocks. We thank Mark Fiers for help with ChIP-seq analysis. We are grateful to Ariel Lindner and Dusan Misevic for discussions and critical feedback on the manuscript. We acknowledge the Nucleomics core facility, Vlaams Instituut voor Biotechnologie (VIB; Flanders Institute for Biotechnology) for their sequencing services, and the scientific and technical assistance of the ICM.Quant imaging core facility (“Investissements d’avenir” program [ANR-10-IAIHU-06] and [ANR-11-INbS-0011]). We thank Best Gene, Inc.; Genetic Services, Inc.; and GenetiVision for the transgenesis services. This work was supported by the program “Investissements d’avenir” ANR-10-IAIHU-06, ICM, VIB, the WiBrain Interuniversity Attraction Pole network (Belspo), and the Paul G. Allen Frontiers Group. R.K.E. was supported by the EMBO long-term fellowship (EMBO ALTF 1056-2011) and the omics@vib fellowship (FP7-PEOPLE-2010-COFUND). N.M. was supported by the Fonds Wetenschappelijke Onderzoeks (FWO) fellowship. B.A.H is an Allen Distinguished Investigator and an Einstein Fellow of the Berlin Institute of Health.

## Author contributions

Conceived and designed the experiments: R.K.E., B.A.H. Bioinformatics: R.K.E. Cloning and fly husbandry: R.K.E., G.H., N.D., Sample preparation and imaging: R.K.E., G.H., S.F.T., A.W.C. ChIP-seq and analysis: N.M. Image analysis: R.K.E., G.H. Data analysis and software: R.K.E. Manuscript preparation: R.K.E., G.H., B.A.H.

## Methods

### Putative target gene predictions and construct design

The list of putative Ato target genes was generated using i-cisTarget webtool^22^ (https://med.kuleuven.be/lcb/i-cisTarget). We used the same lists of up-/downregulated genes in Ato gain or loss of function, as in the original cisTargetX (predecessor of i-cisTarget) publication^17^. For each putative target gene, a suitable fosmid or BAC has been found using the TransGeneOmics database^23,42^ (https://transgeneome.mpi-cbg.de). Recombineering primers were automatically designed using the same database. Fosmid clones were preferred over BACs. Clones with more upstream than downstream sequence and shorter clones were prioritized. The constructs for Ato[mCherry] knock-in were designed in CLC Main Workbench (https://www.qiagenbioinformatics.com/products/clc-main-workbench). The Sanger sequencing assembly and analysis, as well as primer design for sample validation, was also performed in CLC Main Workbench. Annotated BAC and plasmid sequences are available in the project GitHub repository (https://github.com/rejsmont/rdn-wdp-data).

### Plasmids, Fosmids and BACs

Selected fosmids and BACs carrying putative Ato target genes were C-terminally tagged with the TagNG[2xTY1-T2A-Venus-NLS-3xFLAG]^43^ as described in the liquid culture recombineering protocol^44^ (Protocol 8 in the thesis; full text is available online from TU Dresden library https://nbn-resolving.org/urn:nbn:de:bsz:14-qucosa-66452). In short, chloramphenicol (Cm)-resistant bacteria carrying the genomic constructs were transformed with temperature-sensitive (30°C) pRedFlp4 plasmid that contains the homologous recombinase (Red operon) under rhamnose (rha)-inducible promoter and the flippase under anhydrotetracycline (aHT)-inducible promoter. Bacteria were cultured overnight at 30°C with Hygromycin (Hyg) and Cm selection. Fresh cultures were inoculated from the overnight cultures, and the expression of the Red operon was induced with rhamnose. After induction bacteria were transformed with the PCR-amplified recombineering cassette. Recombinants were selected in liquid culture at 30°C under Cm+Hyg+Kan selection. The saturated cultures were used to inoculate an overnight culture on medium with Cm+Hyg+aHT at 30°C to remove the FRT-flanked kanamycin selection cassette. Finally, the last overnight cultures at 37°C with Cm selection were inoculated to remove the helper plasmid. Low salt LB (Sigma L3397) was used as a medium for overnight cultures. Recovery after electroporation was performed in the SOC medium (Sigma S1797). After the last culture bacteria were plated on chloramphenicol (15μg/ml) LB-agar plates and single colonies were selected and verified using a colony PCR with TagT2A_chk_fwd (CGG AGA TGT GGA GGA GAA TC) and TagT2A_chk_fwd (CTT GTC GTC GTC ATC CTT GT) primers. The same primers were used for final construct verification by Sanger sequencing. The integrity of fosmids and BACs was verified by *Xba*I fingerprinting.

The recombinase-mediated cassette exchange construct used to generate the Ato[mCherry] allele (pattB-5’ato-3’ato-attB_ato-mCherry) was based on pattB-5’ato-3’ato-attB_ato-eGFP plasmid (NM / Hassan Lab). mCherry coding sequence was amplified from pTagNG^43^ using XhoI-mCherry-fwd (TGC GCC TCG AGG GCG GAT CTG GCG GAT CTG GCG GAT CTA TGG TGA GCA AGG GCG AGG AGG) and BsiWI-mCherry-rev (GAA TTC ACG TAC GTT ACT TGT ACA GCT CGT CCA TG) primers and cloned into *Xho*I and *Bsi*WI sites of the ato-eGFP plasmid.

### Fly stocks and husbandry

The flies carrying the *Ato[mCherry]* allele were generated using the IMAGO technique^25^. The pattB-5’ato-3’ato-attB_ato-mCherry construct was injected into the ato^w+^ knock-in flies (*vas-phiC31;; ato*^w+^ *16-1/TM6c*, XQ / Hassan Lab). F0 males were crossed to (*w;; TM3 / TM6c*) virgins. Selected recombinants (*w;; Ato[mCherry] / TM3*) were used to establish a homozygous stock used in the subsequent crosses. The 39 tagged fosmids and BACs carrying putative Ato target genes (TG) were used for PhiC37-mediated transgenesis^45^ into the VK00037^46^ landing site flies (*y^1^, M{vas-int.Dm}ZH-2A, w^1118^; PBac{y^1^-attP-3B}VK00037*, Bloomington 24872). Transformants were selected for *3xP3-dsRed* (fosmids) or *w*^+^ (BACs) and balanced with CyO. Male flies (*w/Y; TG-Venus/CyO*) with either *w*^+^ or *3xP3-dsRed* marker were crossed to virgin females (*w; L/CyO; D/TM6C, Sb, Tb*). From this cross, male progenies were selected (*w/Y; TG-Venus/CyO; +/TM6C, Sb, Tb*), and crossed to virgin females (*w; L/CyO-GFP; Ato[mCherry]*). In the last cross, male progenies with the genotype (*w/Y; TG-Venus/CyO-GFP; Ato[mCherry]/TM6C, Sb, Tb*) were selected and crossed to virgin females (*w;; Ato[mCherry]*). Third instar larvae were selected from this cross (*w; TG-Venus/+; Ato[mCherry]*) and used for the imaging experiments. Flies were raised in a temperature and humidity-controlled incubator at 25°C on standard fly food.

### Sample preparation

After selecting the third instar larvae from the imaging cross, the samples were prepared for imaging. Due to the nature of the native fluorescent protein stability and intensity, it was crucial that samples be imaged the same day they were prepared. Larval brain complexes (from ~30 larvae) were dissected under a stereomicroscope. The larvae were dissected in a shallow glass dish with 0.1 M phosphate-buffered saline (PBS), using size 5 forceps and subsequently transferred to a 1.5 ml tube of 0.1 M PBS, kept on ice. The dissections were performed within 30 minutes, followed directly by a fixation with 4% formaldehyde in PBS supplemented with 0.3% TritonX (PBS-T) for 10 minutes. The brain complexes were then rinsed with PBS-T, followed by 3 washes (5, 10, and then 15 minutes). Following the washes, the brains were stained with DAPI (Sigma Aldrich D9564, 1:50,000 in PBS-T) for 15 minutes. Following staining, the brain complexes were washed (5 and then 10 minutes) in PBS-T and then stored in PBS-T at 4°C, for up to 3 hours. Using size 5 forceps and acupuncture needles, the eye-antennal imaginal discs (Figure 4) were dissected from the brain complexes and mounted on a charged slide (~20 discs). The tissue was covered with a thin layer of mounting medium (2.5% DABCO in Mowiol^®^ 4-88) and sealed under a #1.5 coverslip with Fixogum (Marabu).

### Imaging

Individual eye discs were scanned at single-cell resolution with an upright confocal microscope (Olympus FV1200) using a 40x 1.3 NA oil immersion lens. Venus fluorescent signal, which is coupled to TG expression, was captured using a 515 nm diode-pumped solid-state (DPSS) laser (power 0.2μW; AOTF 65%). mCherry fluorescent signal, which is coupled to Ato expression, was captured using a 559 nm DPSS laser (power 1.3μW; AOTF 65%). Venus and mCherry fluorescent signals were imaged using a z-step of 0.3 μm, an image size of 1024 x 512 pixels, and with a voxel size of 0.3 x 0.3 x 0.3 μm. GaAsP photomultipliers were used to detect Venus and mCherry fluorescent signal. DAPI fluorescence signal was captured using a 405 nm diode laser (power 1.3μW; AOTF 2.5%). DAPI fluorescence signal was imaged using a z-step of 0.15 μm, an image size of 2048 x 1024 pixels, and a voxel size (0.15×0.15×0.15 μm). Conventional photomultiplier was used to detect DAPI fluorescent signal. The DAPI channel was oversampled compared with the Venus and mCherry channels because our segmentation algorithm requires a higher resolution to distinguish between individual nuclei. The pixel dwell time was 10μs for all channels. Each channel was scanned separately. Each eye disc took approximately 3 hours to scan, 5-10 discs were scanned per session, and longer wavelength channels were imaged first (mCherry, Venus and DAPI) to reduce photodamage and photobleaching.

### Segmentation

The first step in the applied image segmentation processes was pixel classification using the random forest machine learning approach. Nuclei and background were manually marked using 2 labels (nucleus and background) in the DAPI channel for one z-stack image (randomly selected 512×512 pixels window cropped from full-size image) from each imaging session using Ilastik^47^. Custom Fiji^48^ plugins and scripts (https://github.com/rejsmont/rdn-wdp) utilizing WEKA^49^ and 3D ImageJ Suite^50^ were developed for training and application of the classifier. The classifier was trained based on several filters (Gaussian blur, Hessian, Derivatives, Laplacian, Structure, Edges, Difference of Gaussian, Mean, Median, Variance) with sigma values between 1-16 pixels applied to the training images. Probability maps generated by applying the classifier to each acquired stack were used for the next steps in the segmentation.

The Difference of Gaussians (DoG) filter was applied to the nuclei probability maps with sigma values of 8 and 5, followed by a local maxima filter with 3 pixel radius. The nuclear binary mask (threshold p>0.2) and seeds from the DoG were used for the 3D watershed segmentation. The results of the segmentation were stored as point clouds in per-sample CSV files, together with imaging and segmentation metadata stored in YAML files. Each sample was given a unique, random 6 alphanumeric character identifier.

### Registration

To enable direct comparisons between different samples, sample registration was performed using a custom python script (`analyze2`) available from the data analysis repository (https://github.com/rejsmont/rdn-wdp-python). First, nuclei were read from the point cloud CSV file. Nuclei that differ from the sample mean by more than 50% were rejected. Coordinates of each nucleus were scaled to a unit of mean nucleus diameter in the sample. Rotation of each sample was automatically adjusted so that the visible edge of the disc is on the left and the anterior of the disc is at the top. The morphogenetic furrow (MF) was identified using the Max of Hessian eigenvalue filter applied to the xyz projection of the mCherry channel onto a 1-unit 2D grid. The intensities of each channel were normalized to the mean intensity of mCherry along the MF. The anterior-posterior (y) coordinates of the nuclei were aligned so that the nuclei laying on the MF line have y coordinate equal to zero. For each nucleus, 26 nearest neighbors were identified using a k-dimensional tree (KD-tree). The prominence of mCherry and Venus signal was calculated as a fraction of the value in the nucleus over the mean value in k-26 neighbors. The normalized and registered values were stored in per-sample CSV files and subsequently combined into a single CSV file. 13 samples with extremely low signal to noise ratios or where the disc morphology was distorted were excluded at this step.

### Cell-type classification

To determine the identity of cells in the MF area (−8>y>8) we decided to cluster these cells based on their position along the A-P axis (y), the expression level of Atonal protein (A), and the prominence of Ato expression (P^A^). Due to the number of cells in our samples (millions), simple linkage analysis is impossible due to huge memory requirements. On the other hand, we found that the clustering approaches designed to deal with large datasets (such as K-means^51^ or DBscan^52^) do not perform well on datasets with smooth transitions between the clusters. As changes of both Ato intensity and prominence are continuous is our data, these approaches failed to correctly identify different stages of R8 specification. Faced with these restrictions, we sought to divide a big problem of clustering millions of cells into multiple smaller ones. We randomly selected sets of 20 samples and performed the linkage analysis, using the Euclidean metric and the Ward variance minimization algorithm^53^, on cells in these samples. To recover clusters from linkage analysis we used the maximum number of clusters as a criterion. We optimized this parameter until the R8 cells were resolved in most samples, and found the optimal number of clusters to be six. The random sample selection and clustering have been repeated 1000 times to maximize sample coverage and variability. To identify similar clusters from all the attempts we used a custom clustering algorithm, similar to the K-means method, but optimized to cluster centroids of a known number of clusters (see `cluster_centroids` method in https://github.com/rejsmont/rdn-wdp-python/blob/master/analysis/clustering.py). In short, first, centroids from a random clustering run are used as seeds. Centroids from subsequent runs are iteratively added and the homologous clusters are identified by the shortest Euclidean distance from the seeds. Centroids of newly formed global clusters are used as seeds in the next iteration. After all the samples have been included in the computation, the computed global centroids become new seeds and the whole process is repeated until the distance between the seeds from current and the previous iteration is smaller than a threshold value (1e-10). Subsequently, we selected cells from all samples that are close (Euclidean distance of 1) to global cluster centroids to train a random forest classifier that we then applied to all cells within the MF area. Cell types corresponding to the clusters were identified as follows, in order, removing the identified cluster from the pool: a cluster with the highest Ato prominence was identified as “R8”, a cluster with the highest Ato expression was identified as “MF High”, a cluster with the highest A-P position was identified as Post-MF, a cluster with the lowest A-P position was identified as Pre-MF, from the remaining clusters, the one with the highest Ato expression was identified as “MF Medium” and the one with the lowest, as “MF Low”. Samples in which we failed to recover cells that belong to all six clusters were removed from further analysis. Finally, as in some samples we saw a number of cells that were misclassified, we refined the cluster assignment using A-P position as a discriminator; for example, cells classified as R8 but close (within 2 cell diameters) to the furrow were reclassified as MF High (see the `cleanup_clusters` method for details).

### Chromatin immunoprecipitation (ChIP)

800 eye-antennal discs were isolated from (*w; ato-GFP*) third-instar larvae. Chromatin was immunoprecipitated using anti-GFP (ab290, Abcam) antibody, purified and sequenced following previously published protocols^54^. Raw data of eye-disc samples were deposited in the Gene Expression Omnibus (GEO) database (http://www.ncbi.nlm.nih.gov/projects/geo/) with the accession number GSE110827.

### Data analysis, visualization, and statistics

Final analysis of the data, statistical analysis, and figure plotting was done in Python. The code is available from the data analysis repository (https://github.com/rejsmont/rdn-wdp-python). All data figures were automatically generated (see the figure plotting script for details https://github.com/rejsmont/rdn-wdp-python/blob/master/analysis/figures_paper.py). The gene expression patterns and prominence patterns (Fig. 3, Supplementary Fig. 3 and 5) were plotted as xyz projection of the respective values onto a 1-unit 2D grid. The mean projection was used for the expression values and the max projection was used for the prominence. The ChIP peak area (Fig. 5c) was computed as a product of the peak length (in bp) and the peak height. Expression fold change of putative Ato targets (Fig. 5d) was computed as a fraction of the mean expression in the between Ato-high (MF-High and R8 classes) and Ato-low cells (MF-Low, Pre-MF, and Post-MF). The standard error of the mean (SEM) has been propagated accordingly. The significance of changes has been tested using a two-sided unequal variance z-test with Bonferroni correction. The correlation between expression fold change and the ChIP peak area (Fig. 5b) was assessed from the Pearson product-moment correlation coefficient. P-values were derived from the linear least-squares regression. The significance of gene expression changes between different cell types (Supplementary Fig. 6) was tested using a two-sided unequal variance z-test with Bonferroni correction. Statistical significance was not computed for data points with mean expression values lower than 0.2. Classification of genes into groups of those that strongly or weakly follow the expression of Atonal, those that do not and those that are not expressed in the furrow area was performed manually, based on the significance of expression level changes and the A-P expression profiles in different cell types.

## Supplementary material

**Supplementary Figure 1.**
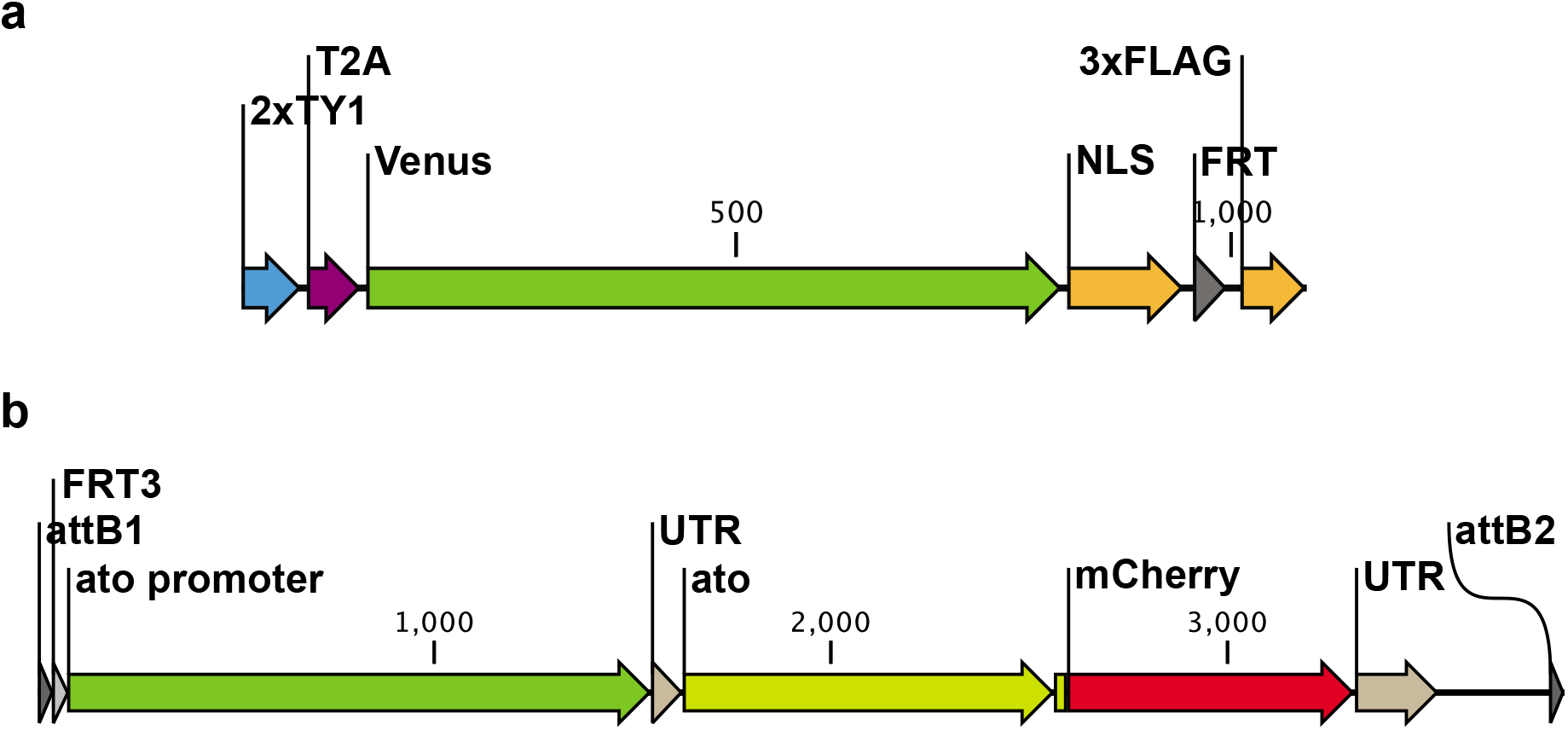
Genetically-encoded fluorescent reporters. (**a**) The transcriptional reporter is designed to be placed C-terminally, just before the target gene STOP codon. The reporter consists of a tandem TY1 epitope, a T2A ribosomal skip sequence, nuclear Venus fluorescent protein, the FRT site that also encodes a degron, and a triple FLAG epitope. (**b**) The ato[mCherry] knock-in construct contains an FRT3 sequence that can be used for mitotic recombination, the *ato* promoter, and the full-length *ato* transcript sequence. The insert is flanked by attB sites used for knock-in using the IMAGO technique.

**Supplementary Figure 2.**
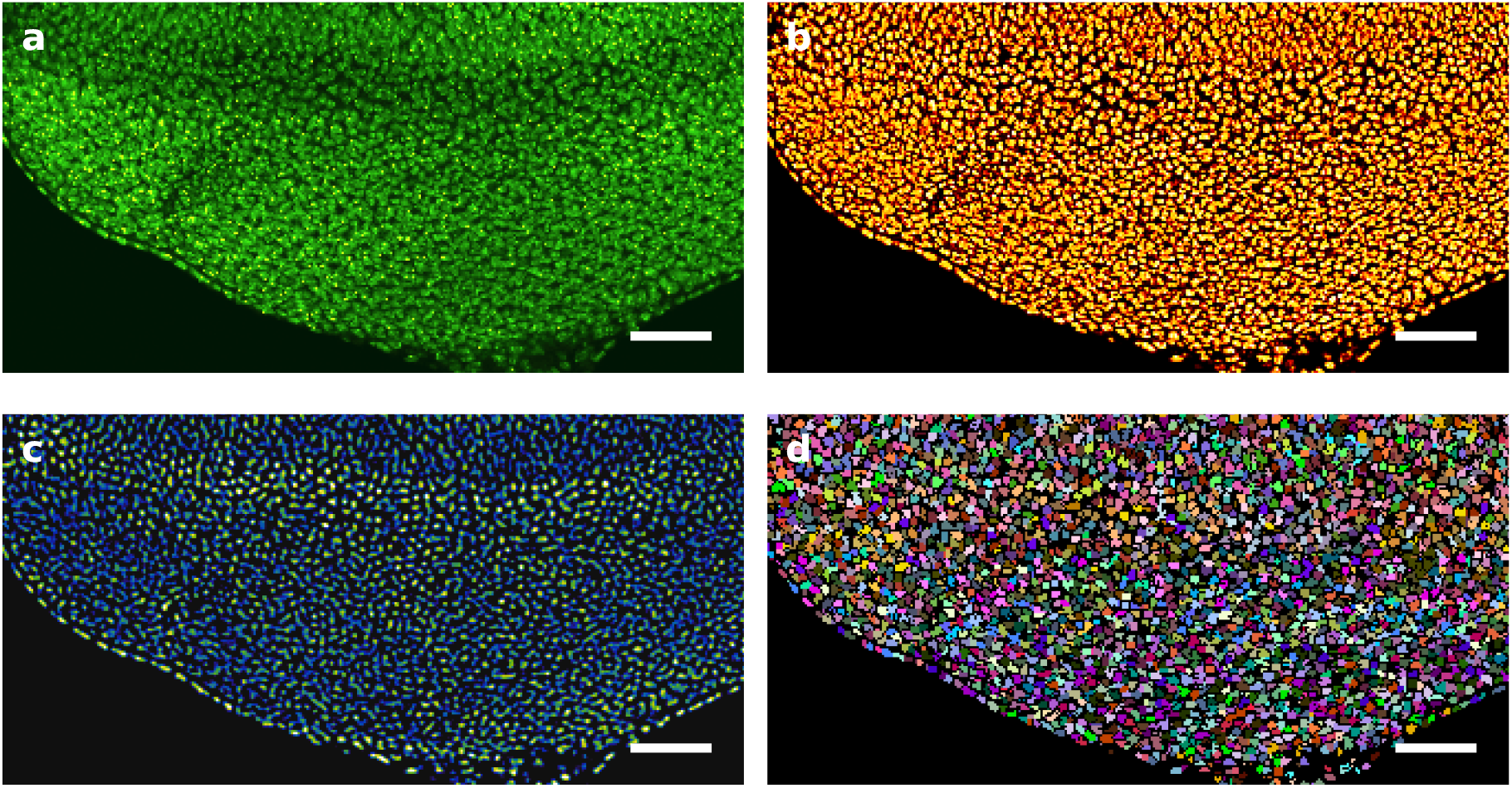
Nuclear segmentation. (**a**) A single section of the DAPI channel used for segmentation (sample 1Q8GA8). Scale bar is 30 μm. (**b**) Pixel classification probability map used to detect the nuclei. (**c**) Seed detection using the difference of Gaussians (DoG) calculated from the probability maps. (**d**) Result of 3D watershed calculated on the probability map using DoG seeds.

**Supplementary Figure 3.**
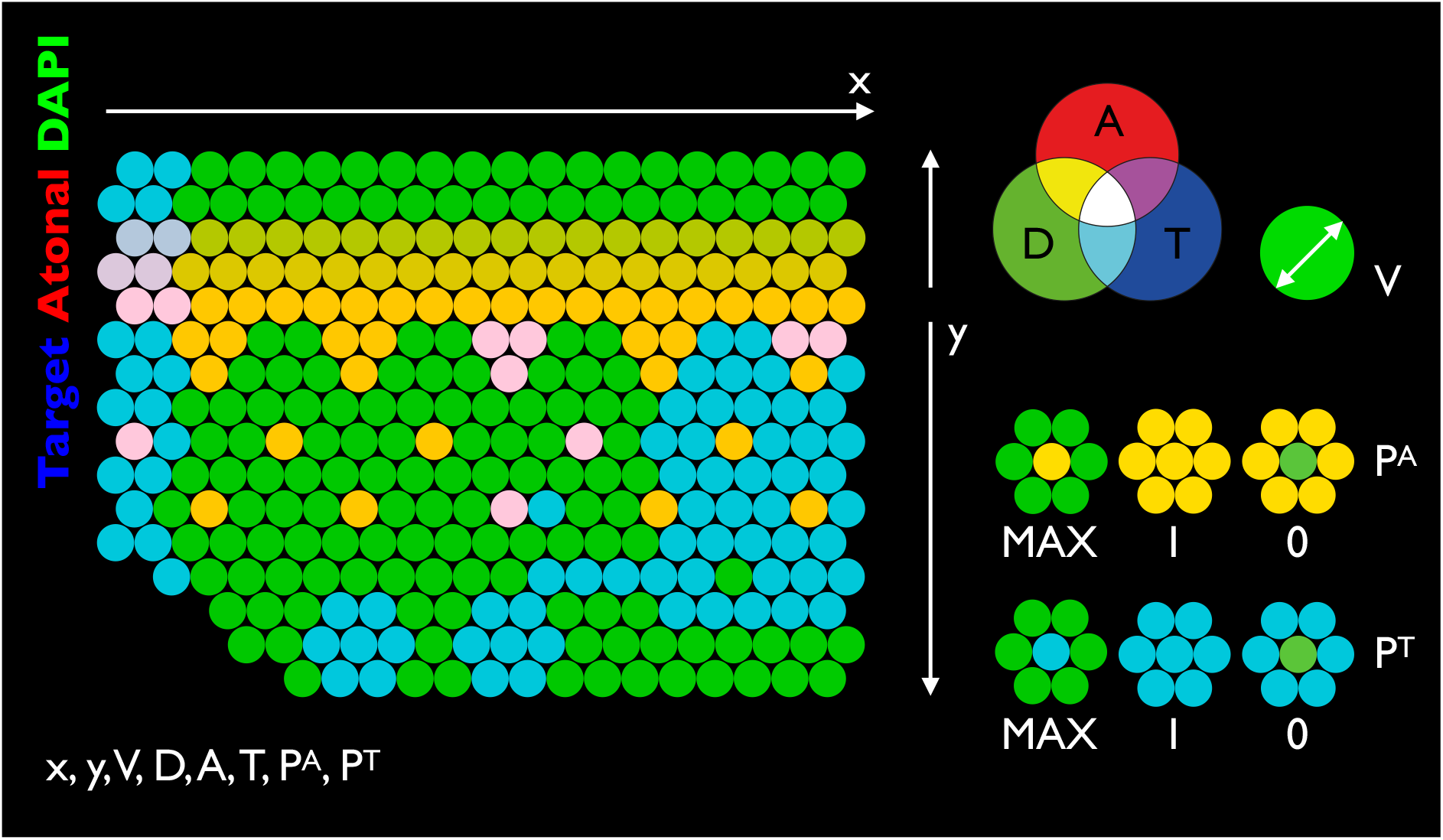
Sample-invariant coordinate system used for sample registration. Each nucleus is represented by eight values. The horizontal (**x**) axis represents the D-V distance of the nucleus from the disc edge. The vertical (**y**) axis represents the A-P distance from the morphogenetic furrow. Both values are expressed in units of mean nuclear diameter in the disc. The nucleus volume (**V**) is expressed in voxels (1 voxel = 3.375×10^−3^ μm^3^). Signal intensities for DAPI (**D**), Ato[mCherry] (**A**) and the T2A-Venus transcriptional reporters (**T**) are normalized to the mean intensity of Ato[mCherry] along the MF. The prominence of Ato[mCherry] (**P^A^**) and the reporter (**P^T^**) is calculated as a mean ratio between the signal intensity in the particular nucleus and its 26 nearest neighbors in 3D space and takes values between 0 (lowest prominence) and infinity (highest prominence). The value of 1 means that the signal intensity in the nucleus is identical to the mean intensity of its 26 neighbors.

**Supplementary Figure 4.**
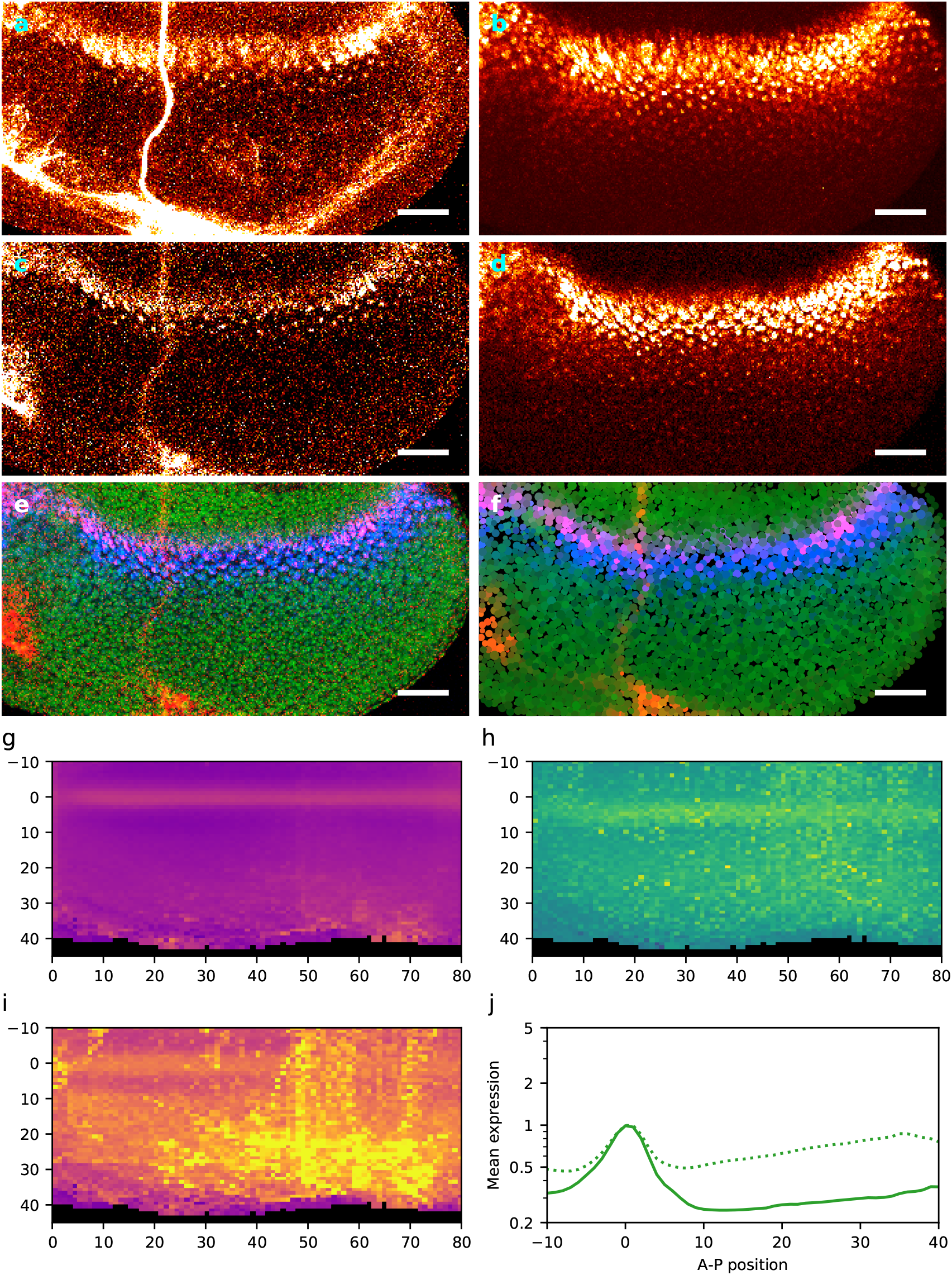
Expression of Atonal in the eye disc. Sample artifacts and the robustness of our approach. (**a**) Maximum intensity z-projection of Ato[mCherry] channel from sample iJbqq8. In addition to Ato[mCherry], the expression 3xP3-dsRed selectable marker is also visible in the posterior part of the disc as well as in the optic nerve. (**b**) Maximum intensity z-projection of ato-T2A-Venus-NLS transcriptional reporter channel from the same sample. (**c**) A single section of the Ato[mCherry] channel. (**d**) A single section of the transcriptional reporter channel. (**e**) A single section composite of all channels (DAPI - green, Ato[mCherry] - red, T2A-Venus - blue). (**f**) Nuclear point cloud from the same section. (**g**) xyz mean intensity projection of normalized Ato[mCherry] signal from all samples. (**h**) xyz maximum projection of Ato[mCherry] prominence in all samples. (**i**) xyz maximum intensity projection of Ato[mCherry] signal from all samples. (**j**) comparison of the A-P profiles of Ato[mCherry] expression measured in BAC (solid line) and all (dotted line) samples.

**Supplementary Figure 5.**
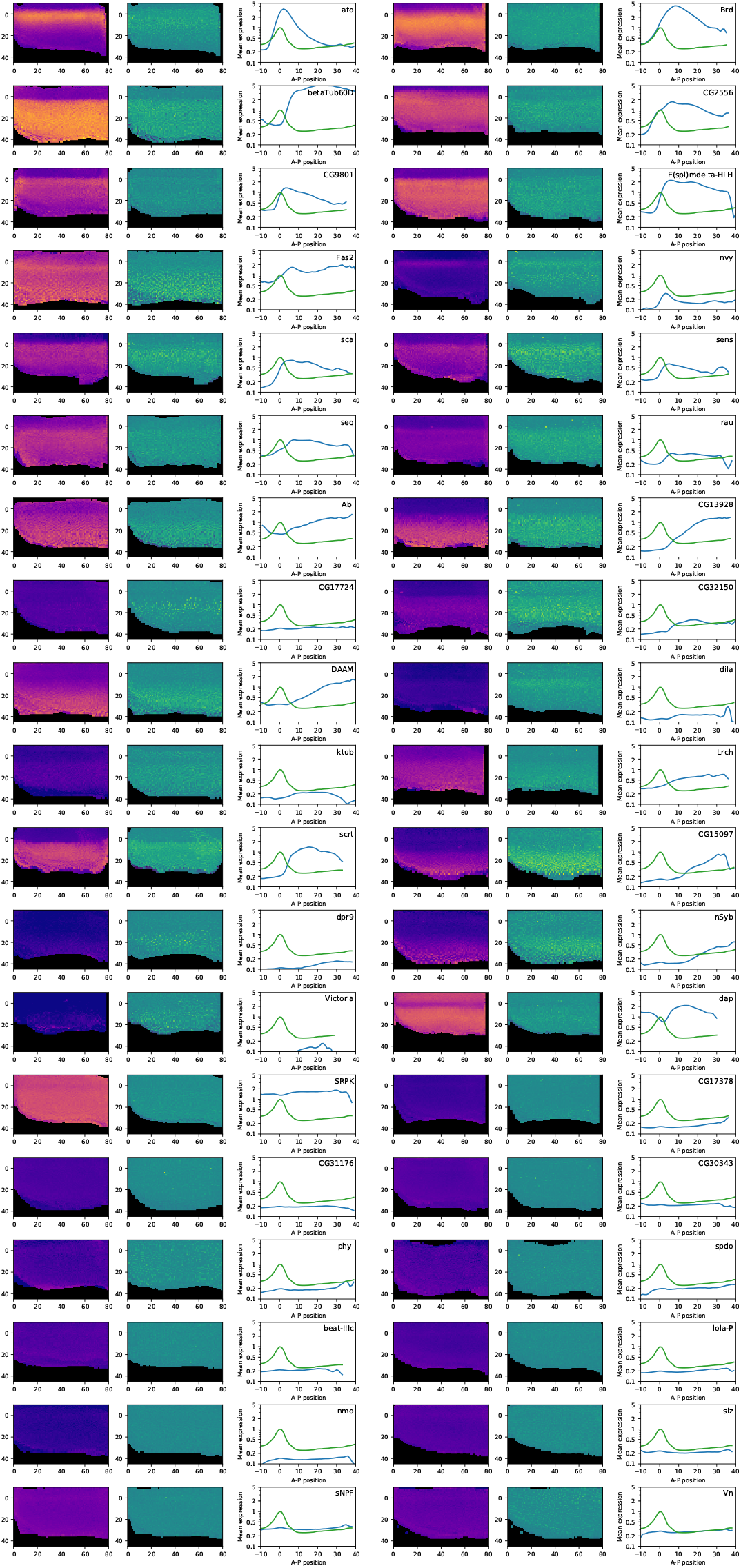
Expression of the putative Ato targets in the eye disc. The left plots show the xyz mean intensity projection of the normalized reporter expression. The middle plots show the xyz maximum projection of the reporter prominence. The right plots show the A-P expression profile of the reporter (blue line) and the Ato protein (green line) as a reference.

**Supplementary Figure 6.**
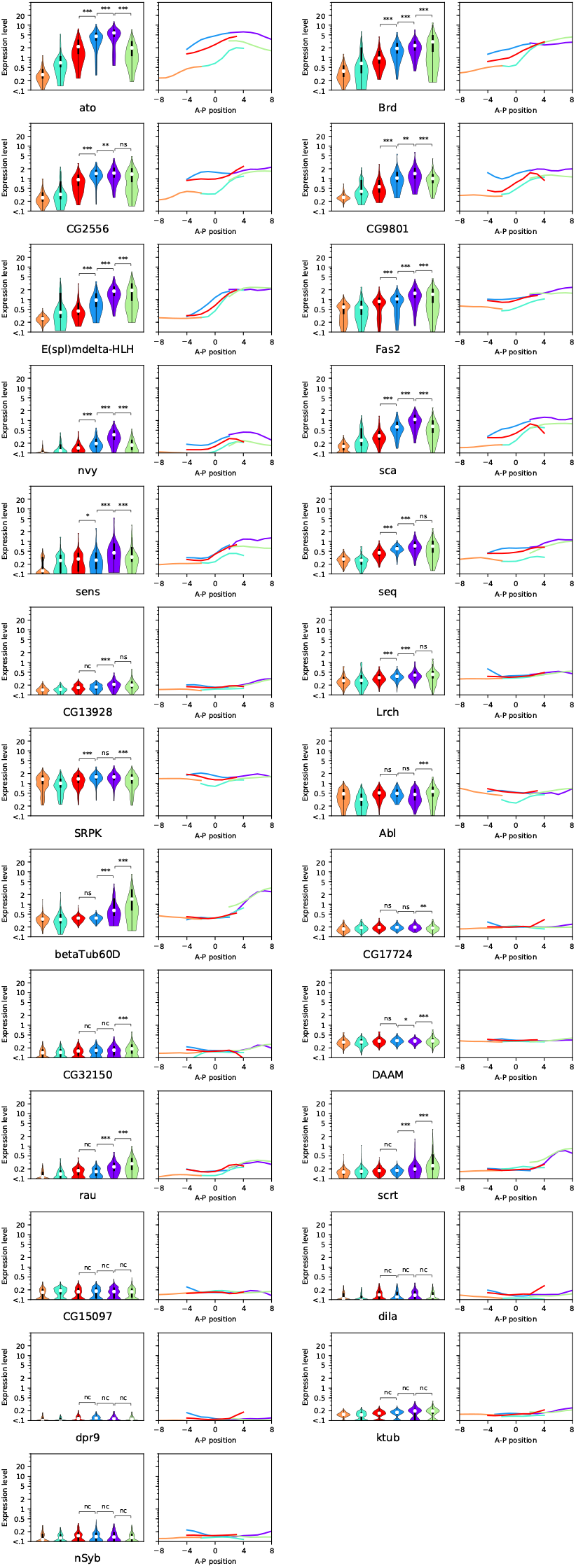
Expression of the putative Ato targets in different cell types. Violin plots show the distribution of the putative target gene expression levels in different cell types in the vicinity of the morphogenetic furrow. Statistical significance of the expression differences in MF-Medium / MF-High, MF-High / R8, and R8 / Post-MF pairs was computed using a two-sample z-test with Bonferroni correction. Line plots show the A-P expression profiles in each cell type. Color code is the same as in figure 4 (violet - R8, blue - MF-High, red - MF-Medium, cyan - MF-Low, orange - Pre-MF, green - Post-MF).

**Supplementary Table 1.**
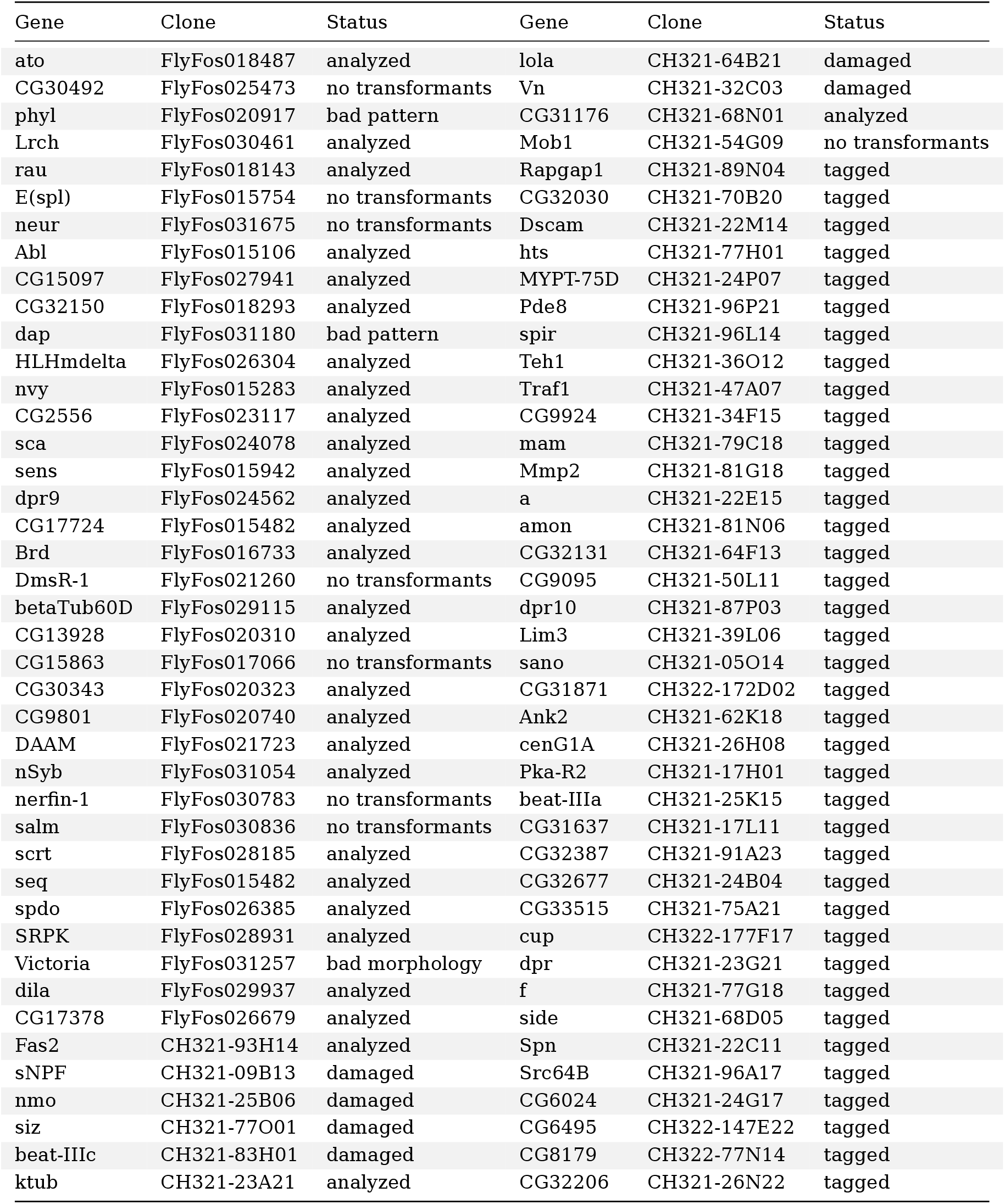
Genes and clones. This table list all predicted targets of Ato, the genomic clones used to create the transcriptional reporters, and the status of their analysis. Clone names are listed according to the FlyFos and p[ACMAN] library naming scheme. Genes that were fully analyzed in this study are labelled “analyzed”. Genes rejected due to bad disc morphology are labelled “bad morphology”. Genes for which the expression pattern was different than published elsewhere are labelled “bad pattern”. Genes that failed to produce transformant flies are labelled “no transformants”. Genes that were not selected for transgenesis at the time of publication are labelled “tagged”.

**Supplementary Table 2.**
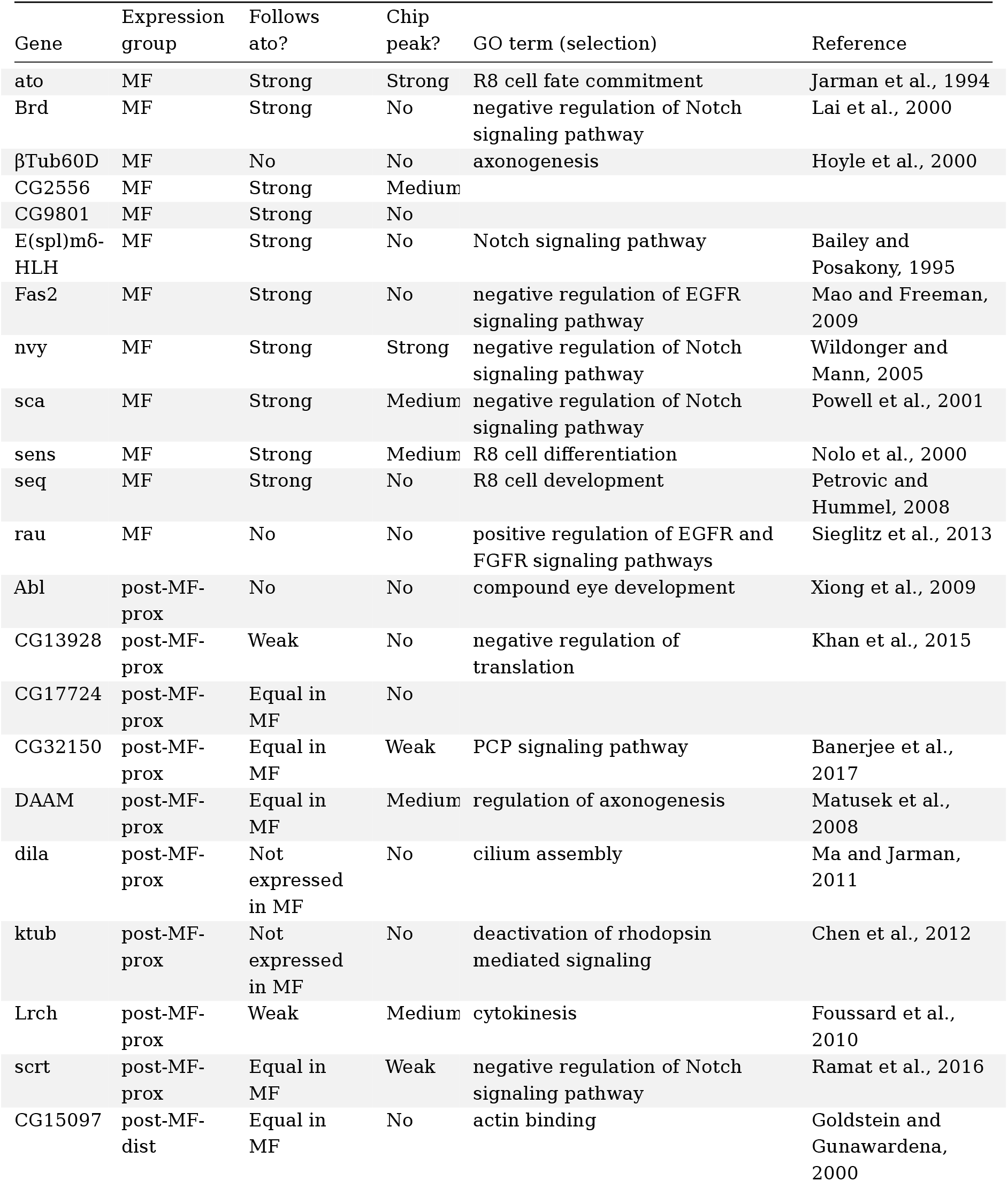
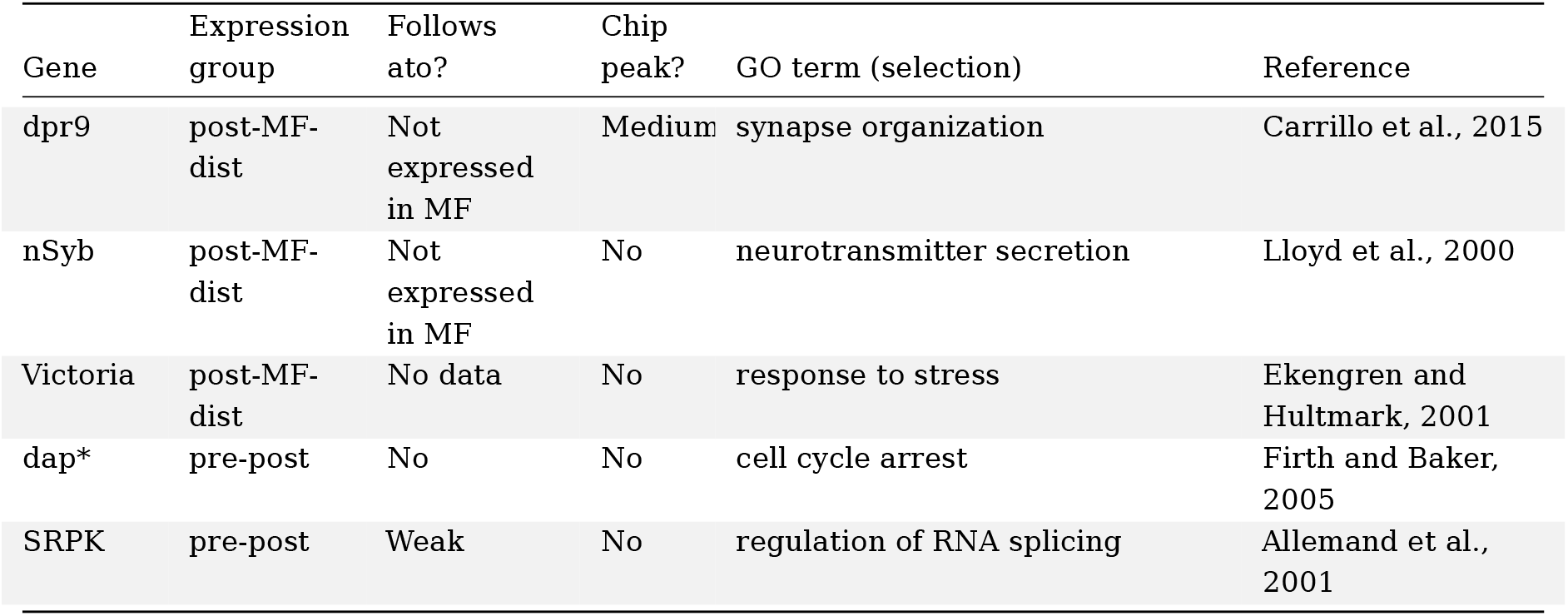
Summary of Ato target gene expression. The list of analysed genes expressed in the eye discs, ordered by the their expression domain. Genes whose expression starts in the morphogenetic furrow (MF) are labelled “MF”. Genes whose expression starts immediately posterior to the MF are labelled post-MF-prox. Genes expressed posterior to the MF are labelled post-MF-dist. Genes expressed both anterior and posterior to the furrow are labelled pre-post. One gene (*dap*) expressed in the disc did not show previously published expression pattern and is marked with an asterisk (*). Expression of the genes was manually assessed in the relationship to Ato. We found genes that showed strong, weak, or no relationship to the Ato expression. Genes that did not change their expression level within the MF region, or that were not expressed there were labelled accordingly. Strength of the ChIP peak was manually assessed as strong, medium or weak. Genes with no detected ChIP signal were labelled accordingly. The Gene Ontology (GO) term relevant to the R8 specification and the supporting reference was is also included in the table.

**Supplementary Table 3.** A-P boundaries of the identified clusters. This table lists the identified cell classes as well as the A-P regions that they occupy. The distances are expressed in the normalized distance units (1 unit is the mean nuclear diameter in a sample) from the morphogenetic furrow. Negative values are anterior to the furrow. The last column shows the color code used to label the cell type in the figures.

| Cluster | Start | End | Color |
| --- | --- | --- | --- |
| Pre-MF |  | -2 | orange |
| MF-low | -1 | 3 | cyan |
| MF-med | -3 | 2 | red |
| MF-high | -1 | 2 | blue |
| R8 | 2 | 6 | violet |
| Post-MF | 2 |  | green |

